# Biofabrication of engineered tissues by 3D bioprinting of tissue specific high cell-density bioinks

**DOI:** 10.1101/2024.09.11.612457

**Authors:** Oju Jeon, Hyoeun Park, J. Kent Leach, Eben Alsberg

## Abstract

Bioprinting of high cell-density bioinks is a promising technique for cellular condensation-based tissue engineering and regeneration medicine. However, it remains difficult to create precisely controlled complex structures and organization of tissues with high cell-density bioink-based bioprinting for tissue specific condensation. In this study, we present newly biofabricated tissues from directly assembled, tissue specific, high cell-density bioinks which have been three-dimensionally printed into a photocrosslinkable and biodegradable hydrogel microparticle supporting bath. Three types of tissue specific high cell-density bioinks have been prepared with individual stem cells or stem cell aggregates by incorporation of growth factor-loaded gelatin microparticles. The bioprinted tissue specific high cell-density bioinks in the photocrosslinked microgel supporting bath condense together and differentiate down tissue-specific lineages to form multi-phase tissues (e.g., osteochondral tissues). By changing the growth factors and cell types, these tissue specific high cell-density bioinks enable engineering of various functional tissues with controlled architecture and organization of cells.

## 1. Introduction

Three-dimensional (3D) bioprinting has gained prominence in the field of tissue engineering and regenerative medicine due to its ability to fabricate complex 3D structures with high resolution and shape fidelity using cell-free or cell-laden bioinks^[1]^. Although 3D bioprinting has brought remarkable capabilities to the field of tissue engineering and regenerative medicine^[2]^, technologies to create and culture individual cell-only-based high-resolution tissues with high shape fidelity that form cell condensations with defined geometries, without an intervening biomaterial scaffold to maintain construct shape and architecture, still face challenges. Previous reports of bioinks composed of only cells either were unable to provide long-term support for 3D printed structures^[3]^ or required use of preformed cell aggregates or strands, which decreases printing resolution, involves extensive preculture time, and/or necessitates expensive equipment ^[4]^.

Recently, we have reported a new strategy enabling the cell condensation-based tissue generation with defined geometries from direct assembly of individual stem cells, which have been 3D bioprinted into a photocrosslinkable shear-thinning and self-healing supporting bath comprised of hydrogel microparticles (microgels)^[5]^. The microgels are prepared by ionic crosslinking of biodegradable and photocurable oxidized and methacrylated alginate (OMA). The OMA microgel supporting bath maintains the individual cell-only bioink 3D printed structure with high resolution and shape fidelity. While the microgel supporting bath allows free movement of a printing needle via its shear-thinning properties, the microgels work as supporting biomaterials for printed cell-only constructs through self-healing properties. Unique to this strategy, after direct 3D bioprinting of the cell-only bioink, subsequent photocrosslinking of the microgels in the supporting bath provides mechanical stability enabling cellular condensation during long-term culture for a defined time-period. The microgel supporting bath can be tuned to fully degrade following cellular condensation formation and/or guided differentiation of 3D printed cells, allowing facile harvesting of the printed constructs without damage. This is in contrast to other microparticle baths that cannot support cell-only printing and condensation formation because they either degrade immediately at 37°C and/or cannot be further crosslinked after printing to provide stabilization during culture^[1c],[3]^.

The purpose of this study was to explore an advanced cell-only 3D bioprinting strategy utilizing tissue-specific high cell-density bioinks by incorporation of tissue-specific growth factor-laden microparticles. The use of microparticle-based growth factor delivery could remove the need for exogenous supplementation of growth factors and allow for increased availability throughout the entire 3D printed constructs and/or in distinct localized regions. Microparticle-based growth factor delivery within the cell-only bioinks enabled sustained release and spatially and temporally controlled presentation of growth factors, promoted localized differentiation of stem cells and enabled formation of multi-tissue constructs. Using this advanced 3D bioprinting platform, multi-phase cellular condensation constructs were created with complex geometries composed of spatially distinct tissue regions.

## 2. Materials and methods

### 2.1. Synthesis of OMA and fabrication of OMA microgel supporting bath

Oxidized alginate (OA) was prepared by reacting sodium alginate with sodium periodate ^[5–6]^. Sodium alginate [10 g, Protanal LF 120M high viscous (251 mPa•S at 1% w/v in water), a generous gift from International Flavors & Fragrances, Inc.] was dissolved in ultrapure deionized water (diH_2_O, 900 ml) overnight. Sodium periodate (0.218 g, Sigma) was dissolved in 100 ml diH_2_O, added to alginate solution under stirring to achieve 2 % theoretical alginate oxidation, and allowed to react in the dark at room temperature for 24 hours. Methacrylation [10 (2OX10MA) and 20 % (2OX20MA) theoretical] was performed to obtain oxidized and methacrylated alginate macromers by reacting OA with 2-aminoethyl methacrylate (AEMA). To synthesize OMA, 2-morpholinoethanesulfonic acid (MES, 19.52 g, Sigma) and NaCl (17.53 g) were directly added to each OA solution (1 L) and then the pH was adjusted to 6.5. N-hydroxysuccinimide (NHS, 0.588 and 1.176 g, Sigma) and 1-ethyl-3-(3-dimethylaminopropyl)-carbodiimide hydrochloride (EDC, 1.944 and 3.888 g, Sigma) were added to the mixture under stirring to activate 10 and 20 % of the carboxylic acid groups of the alginate, respectively. After 5 minutes, AEMA (0.844 and 1.688 g, Polysciences) (molar ratio of NHS:EDC:AEMA = 1:2:1) was added to the solutions, and the reaction was maintained in the dark at room temperature for 24 hours. The reaction mixture was precipitated into excess of acetone, dried in a fume hood, and rehydrated to a 1 % w/v solution in diH_2_O for further purification. The OMAs were purified by dialysis against diH_2_O using a dialysis membrane (MWCO 3500, Spectrum Laboratories Inc.) for 3 days, treated with activated charcoal (5 g/L, 100 mesh, Oakwood Chemical) for 30 minutes, filtered (0.22 μm filter) and lyophilized. To determine the levels of alginate methacrylation, the OMA was dissolved in deuterium oxide (D_2_O, 2 w/v %, Sigma), and ^1^H-NMR spectra were recorded on an NMR spectrometer (600 MHz, Bruker) using 3-(trimethylsilyl)propionic acid-d4 sodium salt (0.05 w/v %, Sigma) as an internal standard. To fabricate OMA microgels, OMA (500 mg) was dissolved in Dulbecco’s modified Eagle’s medium-low glucose (25ml, DMEM-LG, Sigma-Aldrich) containing 0.05 % photoinitiator (PI, 2-Hydroxy-4’-(2-hydroxyethoxy)-2-methylpropiophenone, Sigma) and placed in a 30 ml syringe, and 1 ml of supersaturated CaSO_4_ (0.21g/ml in diH_2_O) was loaded into another 30 ml syringe. After the two syringes were connected with a custom-made female-female lure lock spiral mixing unit (Figure S1), the two solutions were mixed back and forth 40 times, and then they were further mixed back and forth 10 times every 10 minutes for 30 minutes. To visualize the OMA microgels, they were stained with Safranin O (Saf-O) and then imaged using a microscope (Leitz Laborlux S, Leica) equipped with a digital camera (Coolpix 995, Nikon). The mean diameter of the OMA microgels was measured using ImageJ with the images of the Saf-O stained OMA microgels.

### 2.2. Synthesis of methacrylated gelatin and fabrication of gelatin microparticle (GM)

Methacrylated gelatin was synthesized by reaction of type-A gelatin with methacrylic anhydride using a previously described method^[6a]^. Briefly, porcine skin type-A gelatin (10 g, Sigma) was dissolved in 100 ml Dulbecco’s phosphate buffered saline (PBS, Gibco) at 60 °C and stirred until fully dissolved. Methacrylic anhydride (0.5 ml, Sigma, purity ≥ 92 %) was added at a rate of 0.5 ml/minute to the gelatin solution under stirring at 50 °C, and the reaction was maintained in the dark at room temperature for 3 hours. The reaction mixture was precipitated into excess acetone, dried in a fume hood, and rehydrated to a 10 w/v % solution in diH_2_O. The methacrylated gelatin was purified by dialysis against diH_2_O (MWCO 12-14 kDa) for 7 days at 50 °C to remove salts, unreacted methacrylic anhydride and byproducts, filtered (0.22 μm filter) and lyophilized. To fabricate the GMs, methacrylated gelatin (5 g) was dissolved in DMEM-LG (50 ml) containing 0.05 % PI, placed in a petri dish, photocrosslinked under UV (Omnicure^®^ S1000, EXFO Photonic Solution Inc.) at 20 mW/cm^2^ for 5 minutes, blended using a consumer-grade blender (Osterizer MFG, at “pulse” speed) for 180 sec with 100 ml DMEM-LG, and then lyophilized. For growth factor delivery, GMs were loaded with 400 ng TGF β_1_ (PeproTech, Rocky Hill, NJ) or 4 μg BMP-2 (Genscript, Piscataway, NJ) per mg GMs. Briefly, 10 mg GMs were soaked with 400 μl DMEM in an Eppendorf tube, and then 40 μl of TGF β_1_ (100 ng/μl in 4 mM HCL with 0.1 % BSA) and 40 μl of BMP-2 (1 μg/μl in 4 mM HCL with 0.1 % BSA) were respectively added into the tube. After gentle mixing by pipetting, the tubes were incubated at 4 °C overnight.

### 2.3. Preparation of growth factor (GF)-loaded GM incorporated tissue specific high cell-density bioinks

Bone marrow-derived hMSCs from a single donor were isolated using a Percoll^®^ gradient (Sigma Aldrich, St. Louis, MO) and the differential adhesion method ^[7]^. Bone marrow aspirate was obtained from the posterior iliac crest of a donor under a protocol approved by the University Hospital of Cleveland Institutional Review Board. The bone marrow aspirate was washed with DMEM-LG containing 10% prescreened fetal bovine serum (FBS, Sigma-Aldrich), 100 U/ml penicillin and 100 μg/ml streptomycin (1 % P/S, BioWhittaker, Suwanee, GA). Mononuclear cells were isolated by centrifugation in a Percoll^®^ density gradient, and then isolated mononuclear cells were plated at 1.8×10^5^ cells/cm^2^ in DMEM-LG containing 10 % FBS and 1 % P/S. After 4 days of culture in an incubator at 37 °C and 5 % CO_2_, non-adherent cells were removed and adherent cells were maintained in DMEM-LG containing 10 % FBS and 1 % P/S with media changes every 3 days. After 14 days of culture, the cells were passaged at a density of 5 x 10^3^ cells/cm^2^ and further expanded with DMEM LG containing 10% FBS, 1 % P/S and 10 ng/ml fibroblast growth factor-2 (FGF-2, R&D Systems). Passage 3 hMSCs were used in this study.

Human adipose tissue-derived stromal cells (hASCs) were generously provided by the Stem Cell Biology Laboratory at the Pennington Biomedical Research Center (Baton Rouge, LA). Cells were isolated following an Institutional Review Board-approved protocol as previously described^[8]^. Fresh human subcutaneous lipoaspirate from patients undergoing elective liposuction surgery was processed by digestion in collagenase Type IA (Sigma Aldrich), and then density centrifugation was used to isolate the stromal vascular fraction. These cells were plated in tissue culture flasks at 3500 cells/cm^2^ in DMEM-F12 with 10% FBS (HyClone), 1 % P/S and cultured in a humidified 37°C, 5% CO_2_ incubator. The adherent cell population was cryopreserved in medium containing 80% FBS, 10% DMEM, and 10% dimethylsulfoxide (DMSO, Sigma-Aldrich). On thawing, cells were expanded by plating at 3500 cells/cm^2^ in DMEM-F12 (Sigma) medium containing 10% FBS, 1 % P/S and 10 ng/ml FGF-2. Passage 3 hASCs were used in this study.

To prepare individual cell-based chondrogenic high cell-density bioinks, hMSCs were mixed with TGF β_1_-loaded GMs (1×10^6^ cells/0.75 mg GMs). For individual cell-based osteogenic high cell-density bioinks, 1×10^6^ hASCs were mixed with 0.75 mg BMP-2-loaded GMs.

To prepare aggregate-based tissue specific high cell-density bioinks, 2 % w/v agarose microwells were fabricated as previously described ^[9]^. hMSCs (5.94×10^5^ cells/ml) were suspended in a basal pellet medium [BPM, high-glucose DMEM with 1% ITS+ Premix (BD Biosciences, San Diego, CA, 37.5 μg/ml ascorbate-2-phosphate (Wako USA), 10^−7^ M dexamethasone (MP Biomedicals, Irvine CA)]. 1 ml suspension was added into each well of a 24-well plate containing the agarose molds (~ 2000 cells/aggregate), centrifuged for 5 minutes at 300 × *g*, and cultured in a humidified incubator at 37°C with 5% CO_2_. After aggregate formation for 2 days, the aggregates were collected from the wells by gentle pipetting to displace the aggregates from the agarose microwells. After removing the media following centrifugation at 200×g for 1 minute, the aggregates were mixed with GF-loaded GMs (500 aggregates/0.75 mg GMs).

To prepare tissue specific aggregate-based high cell-density bioinks, agarose microwells were fabricated as previously described ^[9]^. hMSCs suspended in BPM were mixed with TGF-β_1_-loaded GMs (1×10^6^ cells/0.75 mg GMs). 1ml hMSC/GM suspension was added into each well of a 24-well plate containing the agarose molds (1500 cells/aggregate), centrifuged for 5 minutes at 300 ×g, and cultured in a humidified incubator at 37°C with 5% CO_2_. After aggregate formation for 2 days, the aggregates were collected from the wells by gentle pipetting to displace the aggregates from the agarose microwells. After removing the media by centrifugation at 300×g for 1 minute, the aggregates containing TGF-β_1_-loaded GMs were loaded into a 1-ml syringe.

### 2.4. Rheological properties of GF-loaded GM incorporated tissue specific high cell-density bioinks and OMA microgel slurry

Dynamic rheological examination of the GF-loaded GM incorporated tissue specific high cell-density bioinks and the OMA microgel slurries was performed to evaluate their shear-thinning, self-healing and rheological properties with a Kinexus ultra+ rheometer (Malvern Panalytical). In oscillatory mode, a parallel plate (25 mm diameter) geometry measuring system was employed, and the gap was set to 1 mm. After each GF-loaded GM incorporated tissue specific high cell-density bioink or OMA microgel slurry was placed between the plates, all the tests were started at 25 ± 0.1 °C, and the plate temperature was maintained at 25 °C. Oscillatory frequency sweep (0.01-1.3 Hz at 1 % strain) tests were performed to measure storage moduli (G’), loss moduli (G”) and viscosity to characterize the shear-thinning and mechanically stability characteristics of the GF-loaded GM incorporated tissue specific high cell-density bioinks and the OMA microgels. Oscillatory strain sweep (0.1-100 % strain at 1 Hz) tests were performed to determine the shear-yielding points at which the GF-loaded GM incorporated tissue specific high cell-density bioinks and the OMA microgel slurries behave fluid-like. To demonstrate the self-healing properties, cyclic deformation tests were performed at 100 % strain with recovery at 1 % strain, each for 1 minute at 1 Hz.

### 2.5. 3D printing of tissue specific high cell-density bioinks

Individual cell-based tissue specific (Figure 1A, i), aggregate-based tissue specific (Figure 1A, ii) and tissue specific aggregate-based (Figure 1A, iii) high cell-density bioinks were loaded into 1-ml syringes (Gastight Syringe, Hamilton Company), connected to 0.5-inch 22G stainless steel needles (Hamilton Company) and mounted into the syringe pump extruder on a commercial syringe pump-based 3D bioprinter (BioX™, Cellink). A petri dish was filled with OMA microgel slurry at room temperature to serve as a supporting bath and placed on the building platform. The tip of each needle was positioned at the center and near the bottom of the dish, and the print instructions were sent to the printer using the in-house software in the BioX™ printer. After 3D printing of the bioinks, OMA microgel supporting mediums with 3D printed constructs were stabilized by photocrosslinking under UV (Omnicure^®^ S1000) at 20 mW/cm^2^ for 1 minute. After slurry photocrosslinking, 3D printed constructs in the photocrosslinked OMA microgel slurry were transferred into 6-well tissue culture plates with BPM and placed in a humidified incubator at 37 °C with 5 % CO_2_.

**Figure 1.**
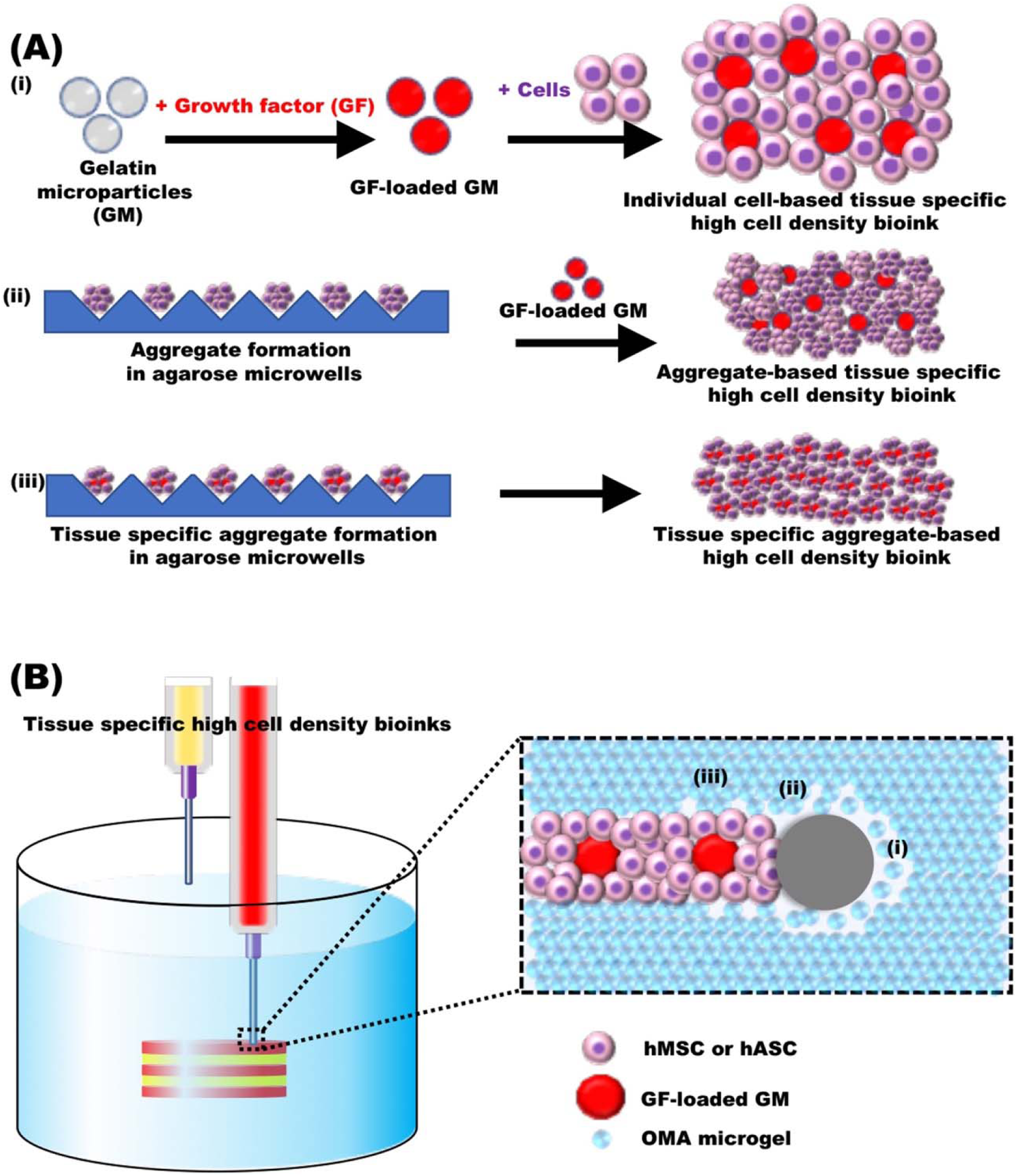
Schematic illustration of **(A)** the tissue specific high cell-density bioink formulations and **(B)** 3D printing of the tissue specific high cell-density bioinks within the OMA microgel supporting bath, which fluidizes via its shear-thinning property. When the printing needle fluidizes the mechanical stable OMA microgels (i, disordered region), tissue specific high cell-density bioinks can fill the shear-thinning region (ii). After the needle passes, the OMA microgel bath can be stabilized by its self-healing property (iii, self-healing region) to firmly hold the printed bioinks^[10]^.

### 2.6. Resolution analysis of 3D printed tissue specific high cell-density bioinks

Linear tissue specific high cell-density bioink filaments were printed in the OMA microgel supporting baths with a 22 needle, the OMA microgel supporting baths were photocrosslinked under UV light at 20 mW/cm^2^ for 1 minute, and then imaged using a microscope (TMS-F, Nikon) equipped with a digital camera (Coolpix 995, Nikon) and fluorescence microscope (ECLIPSE TE 300) equipped with a digital camera (Retiga-SRV). The diameters of the 3D printed hMSC filaments were measured for at least 100 different locations on more than 5 filaments for each bioink using ImageJ (National Institutes of Health).

### 2.7. Osteogenic and/or chondrogenic differentiation of the 3D printed tissue constructs

3D printed constructs in the photocrosslinked OMA microgel supporting baths were osteogenically and/or chondrogenically differentiated by culture with BPM or chondrogenic differentiation media (CPM; BPM + 10 ng/ml TGF-β_1_). The media was changed every other day. After 4 weeks of culture, 3D printed constructs were harvested by simple agitation, fixed in 10 % neutral buffered formalin overnight at 4 °C and stained with Toluidine blue O. Unstained, fixed 3D printed constructs were embedded in paraffin, sectioned at a thickness of 10 µm, stained with Saf-O/Fast Green, Alizarin red S, hematoxylin & eosin (H&E) and Toluidine blue O, and then imaged using a microscope (Leitz Laborlux S, Leica) equipped with a digital camera (Coolpix 995, Nikon). To measure glycosaminoglycan (GAG) production, differentiated 3D printed constructs were homogenized at 35000 rpm for 60 sec using a TH homogenizer (Omni International) in buffer (1 ml, pH 6.5) containing papain (25 μg/ml, Sigma), l-cysteine (2 × 10^−3^M, Sigma), sodium phosphate (50 × 10^−3^M, Thermo Fisher Scientific) and EDTA (2 × 10^−3^M, Thermo Fisher Scientific) and then digested at 65 °C overnight. GAG content was quantified by a 1,9-dimethylmethylene blue assay^[6a]^ and DNA content was measured using the PicoGreen® assay according to the manufacturer’s instructions. GAG content was normalized to DNA content. For quantification of alkaline phosphatase (ALP) activity, DNA content and calcium deposition, differentiated 3D printed constructs were homogenized at 35,000 rpm for 60 sec using a TH homogenizer in 1ml ALP lysis buffer (CelLytic™ M, Sigma). The homogenized solutions were centrifuged at 500 g with a Sorvall Legent RT Plus Centrifuge (Thermo Fisher Scientific). For ALP activity measurements, supernatant (100 μl) was treated with *p*-nitrophenylphosphate ALP substrate (pNPP, 100 μl, Sigma) at 37 °C for 30 minutes, and then 0.1 N NaOH (50 μl) was added to stop the reaction. The absorbance was measured at 405 nm using a plate reader (Molecular Devices). A standard curve was made using the known concentrations of 4-nitrophenol (Sigma). DNA content in supernatant (100 μl) was measured using a Quant-iT PicoGreen assay kit (Invitrogen) according to the manufacturer’s instructions. After an equal volume of 1.2 N HCl was added into each lysate solution, the mixed solutions were centrifuged at 500x*g* with a Sorvall Legent RT Plus Centrifuge. Calcium deposition of the constructs was quantified using a calcium assay kit (Pointe Scientific) according to the manufacturer’s instructions. All ALP activity and calcium deposition measurements were normalized to DNA content.

### 2.8. Statistical analysis

Statistical analyses were performed by one-way analysis of variance (ANOVA) with the Tukey significant difference post hot test using the Prism (Graphpad). A value of p<0.05 was considered statistically significant.

## 3. Results and discussion

In this study, we have developed a new strategy for OMA microgel fabrication with a custom-made spiral mixing unit (Figure S1). This method could eliminate our previously utilized blending process^[5]^, which required preservatives (e.g., 70 % ethanol) for long-term storage and extra washing steps to remove preservatives. These steps could cause premature degradation of the OMA microgels. The microgels formed by this new process with 2OX10MA and 2OX20MA exhibited similar diameters and morphology (Figure S2, A-C) from our previous blending approach^[5]^. The OMA microgels fabricated with our custom-made spiral mixing unit had favorable properties (e.g., shear-thinning, shear-yielding, phase-changing by shear strain and self-healing properties, Figure S2D-I) for use as a supporting medium for high cell-density 3D bioprinting (Figure S2, D-I).

To prepare the tissue specific high-cell density bioinks, we first used individual cells and GF-loaded GMs [Figure 1A, (i)], which can release growth factors in a sustained manner^[11]^. We also utilized the cell aggregates supplemented with GF-loaded GMs [Figure 1A, (ii), Aggregate+GM] and the cell and GF-loaded GMs aggregates [Figure 1A, (iii), Aggregate formed with GM] as the tissue specific high cell-density bioinks. Due to the shear-thinning and self-healing properties of the ionically crosslinked alginate microgels, the printing needle could freely move in the microgel bath and 3D printed tissue specific bioinks could retain their shape and form integrated structures after deposition (Figure 1B). The alginate microgel supporting bath could be further stabilized by photocrosslinking of methacrylated groups on the microgels to provide long-term support for the 3D printed constructs, which is required for self-assembly of 3D printed tissue specific high cell-density bioinks into cellular condensations, differentiation and formation of functional tissues. Since the physical properties of the photocrosslinked alginate microgel supporting bath (e.g., mechanics, swelling ratio and biodegradation rate) are controllable, they can be tuned to support differential needs for fabricating various cellular condensation-based tissues.

Shear-thinning, self-healing and low shear-yielding properties are crucial for bioink printability^[12]^. Shear-thinning, low shear-yielding and liquid-like behaviors ensure that bioinks easily flow under low shear stress during extrusion through a narrow printing nozzle^[13]^, while self-healing allows rapid recovery of the bioinks’ mechanics after printing, maintaining the shape and integrity of the printed structures^[14]^. To demonstrate the favorable printing properties of the tissue specific high cell-density bioinks, we evaluated their rheological properties.

All of the tissue specific high cell-density bioinks exhibited shear-thinning behavior as characterized by the decreasing viscosity with increasing shear rate (Figure 2A). In addition, all of the tissue specific high cell-density bioinks exhibited low shear yield stress (Figure 2B-C), indicating that only a small shear stress (< 1 Pa) is required to initiate flow of the tissue specific high cell-density bioinks. Frequency sweep tests of the tissue specific high cell-density bioinks showed significantly higher G’ than G”, indicating that all of the tissue specific high cell-density bioinks are mechanically stable (Figure 2D). In the shear strain sweep test, G’ of the tissue specific high cell-density bioinks rapidly decreased at approximately 7 %, and G” of the tissue specific high cell-density bioinks surpassed G’ at approximately 25 % shear strain (Figure 2E), indicating the phase change from solid-like to liquid-like behavior. Importantly, all of the tissue specific high cell-density bioinks exhibited self-healing when investigated under cyclic strain sweeps by alternating 1 % (low) and 100 % (high) shear strain. They showed consistent responses of shear moduli (G” > G’ crossover) to high strain (100 %) and rapid and repeatable recoveries to their original state at low shear strain (1 %) (Figure 2F). The measured mechanical responses support all of the tissue specific high cell-density bioinks being well-suited for extrusion-based 3D bioprinting.

**Figure 2.**
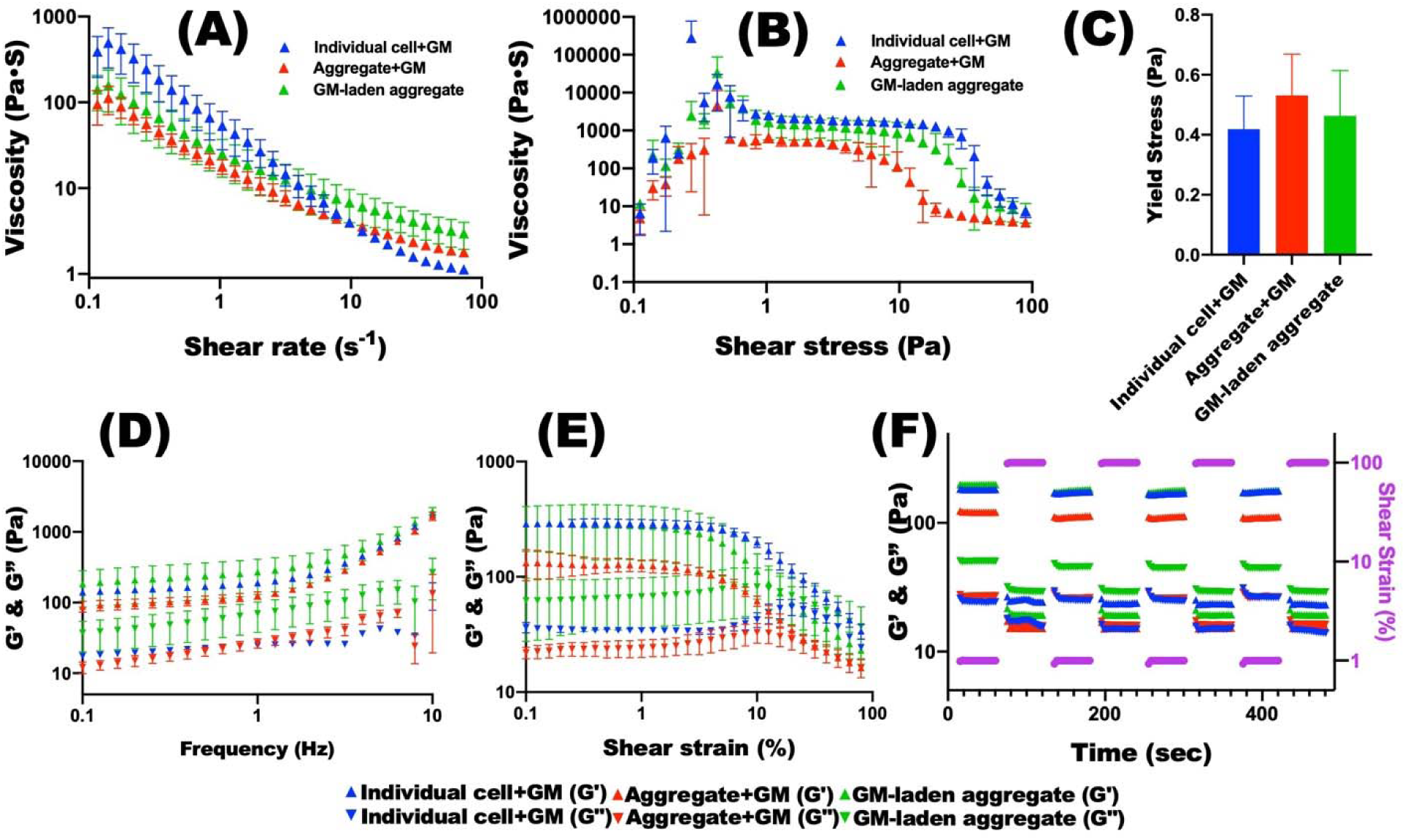
Characterization of the tissue specific high cell-density bioinks. Viscosity measurements of the tissue specific high cell-density bioinks as a function of **(A)** shear rate and **(B)** shear stress demonstrate their shear-thinning and shear-yielding behaviors, respectively (N=3). **(C)** Quantification of the shear-yield stress from shear-stress sweep tests reveals that the tissue specific high cell-density bioinks can flow by low shear-yield stress (N=3). **(D)** Frequency sweep tests indicate that the tissue specific high cell-density bioinks are mechanically stable (G’ > G”) (N=3). **(E)** Strain sweep tests of the tissue specific high cell-density bioinks as a function of shear strain. The G’ and G” crossover indicate their gel-sol transition at approximately 25 % shear strain (N=3). **(F)** Shear moduli changes during dynamic strain tests of the tissue specific high cell-density bioinks with alternating low (1 %) and high (100 %) shear strains at 1 Hz frequency demonstrate their self-healing properties by showing rapid transition between their solid-like and liquid-like behaviors within seconds.

The printed bioinks should maintain high shape fidelity of the 3D printed structures without deformation and flowing after being extruded through the printing needle. To evaluate the retention of the bioink in the microgel supporting bath, microfilaments of the tissue specific high cell-density bioinks were prepared with individual human bone marrow-derived stem cells (hMSCs) or human adipose tissue-derived stromal cells (hASCs) with GF-loaded GMs and printed into the photocrosslinkable alginate microgel supporting baths with different methacrylation degrees of alginate microgels to compare resulting resolutions. Regardless of cell type or alginate degree of methacrylation, all of filaments exhibited similar diameters with high resolution and narrow filament diameter distribution (Figure 3A and B). Therefore, constructs 3D printed into the alginate microgel slurry baths with tissue specific individual hMSC and hASC-based (Figure 3C and D) bioinks exhibited high shape fidelity. After photocrosslinking of the alginate microgel supporting bath, 3D printed constructs with transforming growth factor-beta1 (TGF-β_1_)-loaded GM incorporated hMSC bioink could be cultured in BPM. After 4 weeks of chondrogenic differentiation, successful cartilage tissue formation by cell condensation formation and differentiation of 3D printed hMSCs with delivery of TGF-β_1_ using GMs was confirmed via Saf-O/Fast Green staining (Figure 3E, i and iii) and further quantification of glycosaminoglycan (GAG) production (Figure 3F), which were similar to the positive control [no growth factor loaded gelatin microparticle (Empty GM)/CPM; Figure 3E, ii and iv and Figure 3F]. There were no significant differences of GAG, DNA and GAG/DNA values among four groups (Figure 3F and Figure S3A and B).

**Figure 3.**
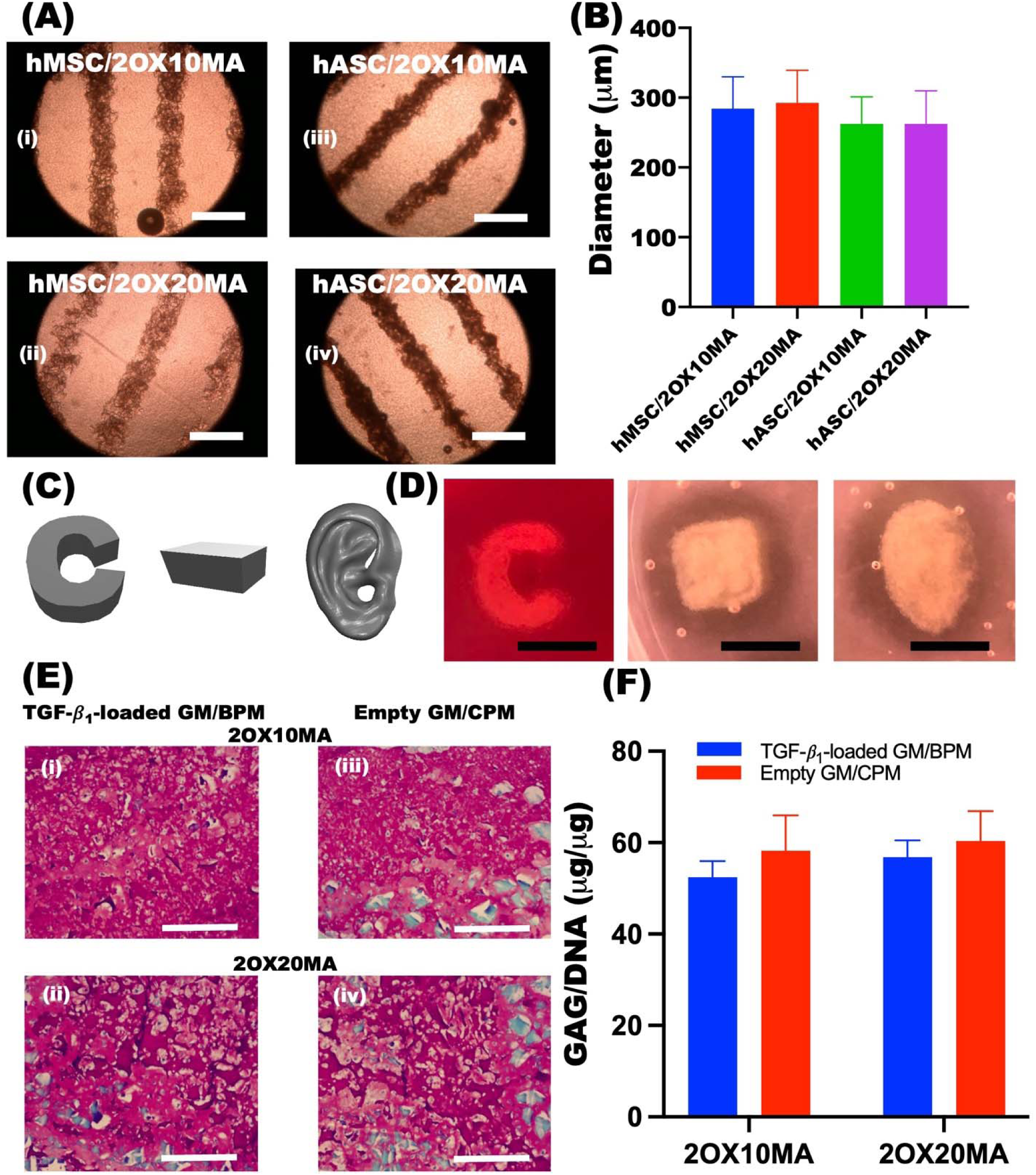
Characterization of resolution and shape fidelity and chondrogenic differentiation of 3D bioprinted constructs. **(A)** Photomicrographs of the 3D printed individual cell-based tissue specific bioink filaments (i and ii; TGF-β_1_-loaded GM + hMSCs. iii and iv; BMP-2-laoded GM + hASCs) into OMA microgel supporting baths with a 22 G printing needle and **(B)** quantification of their mean diameters, demonstrating the capability of high-resolution printing with narrow filament diameter distribution. Scale bars indicate 500 μm. **(C)** Digital images and **(D)** photographs of the 3D printed structures, demonstrating high shape fidelity. The scale bars indicate 5 mm. **(E)** Photomicrographs of Saf-O/Fast Green stained construct sections cultured in BPM (**i** and **ii**) and CPM (**iii** and **iv**). The scale bars indicate 500 μm. **(F)** Quantification of GAG/DNA in the chondrogenically differentiated 3D printed constructs. These demonstrate hMSC differentiation and deposition of chondrogenic ECM in the individual cell-based tissue specific bioink printed constructs (N=3).

Since the GMs allow easy loading and controlled release of various tissue specific signaling biomolecules, multi-phase tissue formation in a defined region could be achieved using the individual cell-based tissue specific high cell-density bioinks. To demonstrate the possibility of multi-phase tissue formation by the 3D printed individual cell-based tissue specific bioinks, tubular structured osteochondral construct comprised of bone- and cartilage-like tissue were bioprinted using osteogenic bioink and chondrogenic bioink in an alternating fashion with a 2 mm thickness for each alternating layer into the photocrosslinkable alginate microgel supporting bath (Figure 4A) according to a digital image Figure S7A). After 4 weeks of osteogenic and chondrogenic differentiation in BPM, the 3D printed construct maintained the initial 3D printed structure and had precisely assembled into multi-cell phenotypes and a multiphase tissue (Figure 4B-I). Individual hMSC-based chondrogenic bioink formed cartilage-like tissue (Figure 4D and E) and individual hASC-based osteogenic bioink formed bone-like tissue (Figure 4F and G). The chondrogenic phase was strongly stained with Saf-O (Figure 4D) while the osteogenic phase was mainly stained with Fast green (Figure 4E). In contrast, the chondrogenic phase was weakly stained with Alizarin red S (Figure 4F) while the osteogenic phase was strongly stained with Alizarin red S (Figure 4G). Importantly, a clear chondrogenic and osteogenic interface was observed between the cartilage and bone phases of the construct by H&E (Figure 4H) and Alizarin red S (Figure 4I) staining. Quantification of GAG content, alkaline phosphatase (ALP) activity and calcium deposition of separate tissue phases supported the histological findings in the chondrogenically (Chondral) and osteogenically (Osteo) differentiated tissue constructs (Figure 4J and K). Significantly higher GAG (Figure S4A) and GAG/DNA (Figure 4J) were found in the chondrogenic phase (Chondral) compared to the osteogenic phase (Osteo), while significantly higher ALP (Figure S4B), ALP/DNA (Figure 4J), calcium (Figure S4C) and calcium/DNA (Figure 4K) were found in the osteogenic phase (Osteo) compared to the chondrogenic phase (Chondral). There was no significant difference in DNA between the two phases (Figure S4D). These results demonstrated successful biofabrication of multiphase osteochondral tissue constructs by 3D bioprinting of TGF-β_1_ and BMP-2-loaded GMs with individual cell-based bioinks.

**Figure 4.**
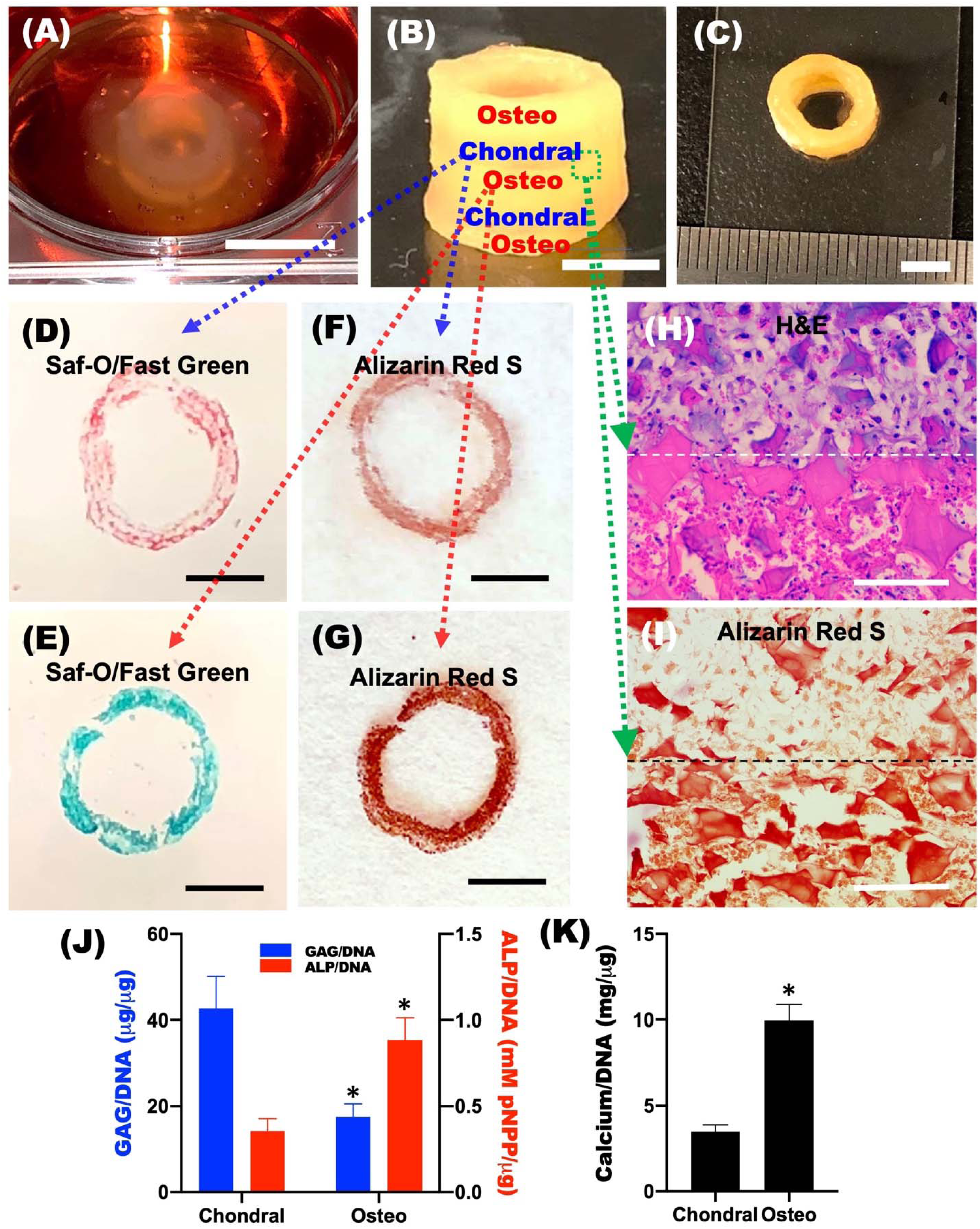
**(A)** 3D printed multi-cell phenotype tube in the photocrosslinkable alginate microgel slurry bath using BMP-2-loaded GM-incorporated hASC bioink and TGF-β_1_-loaded GM-incorporated hMSC bioink. Scale bar indicates 10 mm. **(B)** Side and **(C)** top photographs of 3D printed osteochondral tissue construct after 4 weeks of osteogenic and chondrogenic differentiations in BPM. Scale bars indicate 5 mm. Photomicrographs of Safranin-O/Fast green stained **(D)** chondrogenic and **(E)** osteogenic phases and Alizarin Red S stained **(F)** chondrogenic and **(G)** osteogenic phases of the multi-tissue type osteochondral construct sections. Scale bars indicate 5 mm. Photomicrographs of **(H)** H&E and **(I)** Alizarin Red S stained osteochondral interface of the osteochondral construct sections. Osteo: osteogenic phase; Chondral: chondrogenic phase. Scale bars indicate 500 μm. **(J)** GAG/DNA and ALP/DNA and (K) calcium/DNA contents of chondrogenic and osteogenic phases of the osteochondral construct (N=3). *p<0.05 compared to chondrogenic phase.

Since multicellular aggregates facilitate a high density of cell-cell interactions and self-assemble by fusing with each other, which are valuable for functional and scalable tissue formation, they have been recently used as bioinks and supporting baths for 3D bioprinting applications^[5, 15]^. However, it is still difficult to precisely control the architectures of cell aggregates to mimic sophisticated 3D structures of natural tissues with high resolution and fidelity. In this study, tissue specific aggregate-based bioinks were prepared by incorporation of GF-loaded GMs for specific tissue formation (Figure 1A, ii-iii). To demonstrate if our individual cell printing platform is also suitable for 3D printing of aggregate-based bioinks with high resolution and shape fidelity, hMSC aggregates were prepared using a custom-made agarose mold (Figure S5 and 6). These aggregate-based bioink filaments were then 3D printed into the photocrosslinkable alginate microgel supporting baths. Both Aggregate+GM (Figure 5A) and Aggregate formed with GM (Figure 5B) bioinks exhibited similar resolution compared to the individual cell-based bioink (Figure 3A). The 3D printed structures [letter ‘C’ and tubes (Figure S7)] with the Aggregate+TGF-β_1_-loaded GM (Figure 5D) and Aggregate formed with TGF-β_1_-loaded GM (Figures 5E) bioinks exhibited high shape fidelity compared to their digital image (Figure 5C). After 3D printing of these tissue-specific aggregate-based bioinks, photocrosslinking of the alginate microgel supporting baths could provide long-term support for 3D printed constructs, and the high shape fidelity was well maintained during culture (Figure 5F and G). After 4 weeks of chondrogenic differentiation in BPM, successful cartilage tissue formation was confirmed via Saf-O/Fast green staining (Figure 5H, i and ii) and further quantification of glycosaminoglycan (GAG) production (Figure 5I), which were similar to the positive control [Empty GM (no GF-loaded GM) and cultured in CPM; Figure 5H, iii and iv]. There were no significant differences of GAG, DNA and GAG/DNA values between the four groups (Figure 5I and Figure S8).

**Figure 5.**
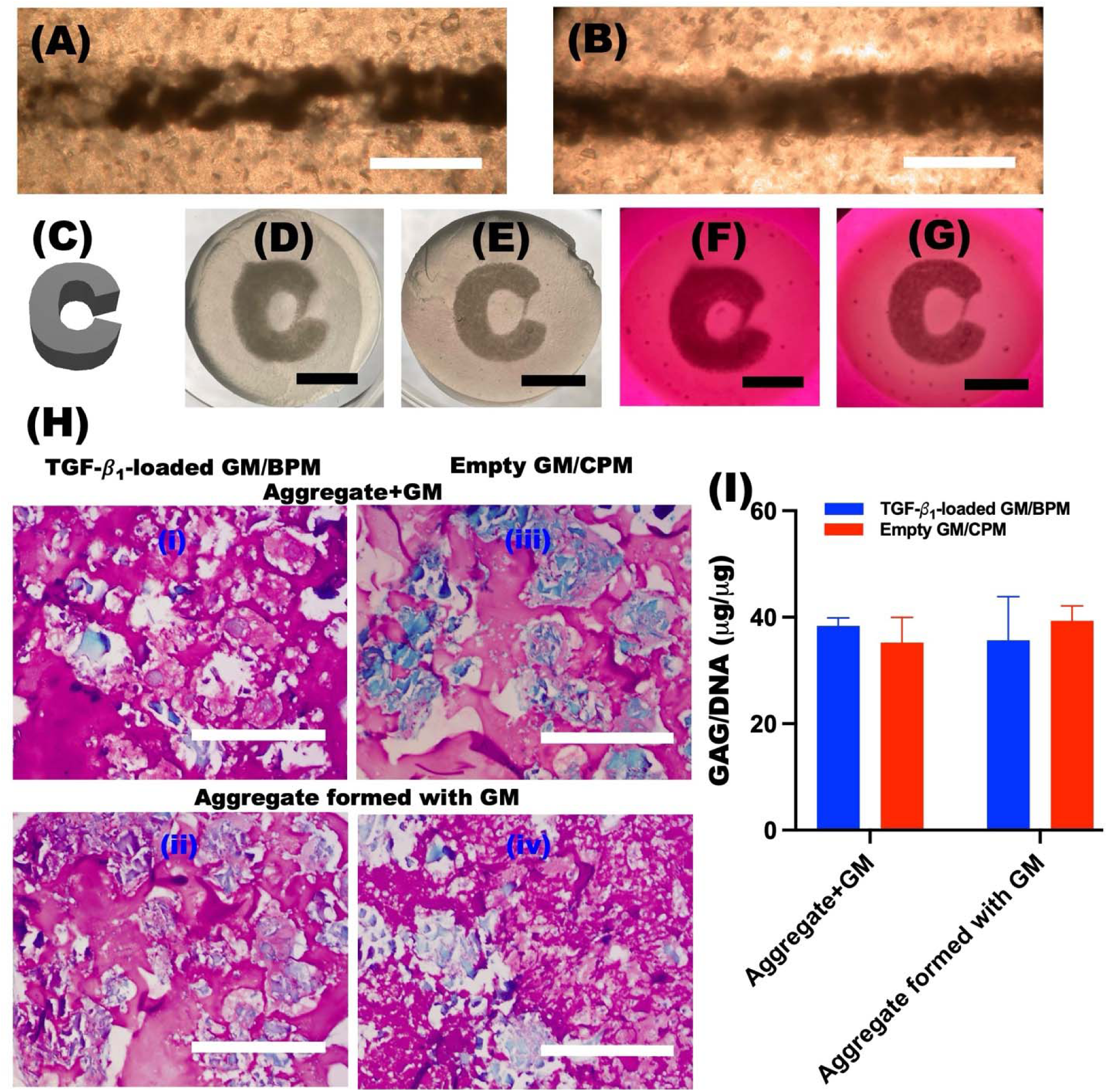
Photomicrographs of 3D bioprinted filaments of **(A)** the Aggregate+GM and **(B)** Aggregate formed with GM bioinks with a 22 G printing needle. Scale bars indicate 500 μm. **(C)** Digital image and photographs of letter “C” using **(D)** the Aggregate+GM and **(E)** Aggregate formed with GM bioinks into OMA microgel supporting bath using a 22 G printing needle after photocrosslinking. Photographs of chondrogenically differentiated 3D printed letter “C” using **(F)** the Aggregate+GM and **(G)** Aggregate formed with GM bioinks after 4 weeks culture in BPM. **(H)** Photomicrographs of Saf-O/Fast Green-stained construct sections cultured in BPM (**i** and **ii**) and CPM (**iii** and **iv**) and **(I)** quantification of their GAG/DNA contents (N=3). The scale bars indicate 500 μm.

Native tissues and organs normally consist of multiple cell types and extracellular matrix (ECM) which are integrated in micro- and/or macroscale spatial patterns^[16]^. Spatiotemporal presentation of multiple biochemical and physical signals is critical for localized cell differentiation and maturation to engineer complex tissues. Therefore, our tissue specific bioink system offers great potential for investigation of the role of spatial presentation of bioactive signals. By 3D bioprinting of multiple tissue specific high cell-density bioinks, we can precisely control the formation of complex multi-tissue structures.

Although there has been a great deal of interest in delivery of bioactive molecules within 3D bioprinted constructs, to the best of our knowledge, our 3D printing platform is the first report of a system capable of spatial presentation of tissue-specific growth factors within high cell-density bioinks, eliciting location specific control over cellular function, such as specific stem cell differentiation into multi-cell phenotypes and multiphase tissue formation with high shape fidelity. Therefore, our approach may open new avenues for the precision engineering of complex tissues using high cell-density bioinks, providing unprecedented control over multicellular function and tissue development.

## Acknowledgements

The authors gratefully acknowledge funding support from the Department of Veterans Affairs, Veterans Health Administration, Office of Research and Development, Rehabilitation Research and Development Service under award number RX004288 and the National Institutes of Health’s National Institute of Arthritis and Musculoskeletal and Skin Diseases under award number R01AR081448. The contents of this publication are solely the responsibility of the authors and do not necessarily represent the official views of the Department of Veterans Affairs or the National Institutes of Health.

## Data availability

The dataset generated and/or analyzed during the current study are available from the corresponding authors upon request.

## References

[1] aC. Mandrycky, Z. Wang, K. Kim, D. H. Kim, Biotechnol Adv 2016, 34, 422; bK. H. Song, C. B. Highley, A. Rouff, J. A. Burdick, Advanced Functional Materials 2018, 28; cT. J. Hinton, Q. Jallerat, R. N. Palchesko, J. H. Park, M. S. Grodzicki, H. J. Shue, M. H. Ramadan, A. R. Hudson, A. W. Feinberg, Sci Adv 2015, 1, e1500758; dX. Zhou, T. Esworthy, S. J. Lee, S. Miao, H. Cui, M. Plesiniak, H. Fenniri, T. Webster, R. D. Rao, L. G. Zhang, Nanomedicine 2019, 19, 58; eN. J. Castro, J. O’Brien, L. G. Zhang, Nanoscale 2015, 7, 14010; fY. Shanjani, C. C. Pan, L. Elomaa, Y. Yang, Biofabrication 2015, 7, 045008; gT. Anada, C. C. Pan, A. M. Stahl, S. Mori, J. Fukuda, O. Suzuki, Y. Yang, Int J Mol Sci 2019, 20; hN. Sears, P. Dhavalikar, M. Whitely, E. Cosgriff-Hernandez, Biofabrication 2017, 9, 025020.

[2] N. A. Sears, D. R. Seshadri, P. S. Dhavalikar, E. Cosgriff-Hernandez, Tissue Eng Part B Rev 2016, 22, 298.

[3] T. Bhattacharjee, C. J. Gil, S. L. Marshall, J. M. Uruena, C. S. O’Bryan, M. Carstens, B. Keselowsky, G. D. Palmer, S. Ghivizzani, C. P. Gibbs, W. G. Sawyer, T. E. Angelini, Acs Biomaterials Science & Engineering 2016, 2, 1787.

[4] aY. Yu, K. K. Moncal, J. Q. Li, W. J. Peng, I. Rivero, J. A. Martin, I. T. Ozbolat, Scientific Reports 2016, 6; bC. S. Ong, T. Fukunishi, H. T. Zhang, C. Y. Huang, A. Nashed, A. Blazeski, D. DiSilvestre, L. Vricella, J. Conte, L. Tung, G. F. Tomaselli, N. Hibino, Scientific Reports 2017, 7.

[5] O. Jeon, Y. B. Lee, H. Jeong, S. J. Lee, D. Wells, E. Alsberg, Mater Horiz 2019, 6, 1625.

[6] aO. Jeon, D. W. Wolfson, E. Alsberg, Adv Mater 2015, 27, 2216; bO. Jeon, D. S. Alt, S. M. Ahmed, E. Alsberg, Biomaterials 2012, 33, 3503.

[7] aS. E. Haynesworth, J. Goshima, V. M. Goldberg, A. I. Caplan, Bone 1992, 13, 81; bD. P. Lennon, S. E. Haynesworth, S. P. Bruder, N. Jaiswal, A. I. Caplan, In Vitro Cellular & Developmental Biology-Animal 1996, 32, 602.

[8] S. G. Dubois, E. Z. Floyd, S. Zvonic, G. Kilroy, X. Wu, S. Carling, Y. D. Halvorsen, E. Ravussin, J. M. Gimble, Methods Mol Biol 2008, 449, 69.

[9] C. E. Vorwald, S. S. Ho, J. Whitehead, J. K. Leach, Methods Mol Biol 2018, 1758, 139.

[10] O. Jeon, Y. B. Lee, H. Jeong, S. Lee, D. Wells, E. Alsberg, Materials Horizons 2019.

[11] L. D. Solorio, E. L. Vieregge, C. D. Dhami, P. N. Dang, E. Alsberg, J Control Release 2012, 158, 224.

[12] aA. Schwab, R. Levato, M. D’Este, S. Piluso, D. Eglin, J. Malda, Chemical Reviews 2020, 120, 10850; bX. Chen, A. F. Anvari-Yazdi, X. Duan, A. Zimmerling, R. Gharraei, N. K. Sharma, S. Sweilem, L. Ning, Bioactive Materials 2023, 28, 511; cC. J. Gao, L. Tang, H. W. Qu, M. M. Wu, T. Zhou, C. Y. Wen, P. P. Wang, N. Xu, C. S. Ruan, Advanced Functional Materials 2024, 34; dO. Jeon, Y. B. Lee, S. J. Lee, N. Guliyeva, J. Lee, E. Alsberg, Bioactive Materials 2022, 15, 185.

[13] aJ. Emmermacher, D. Spura, J. Cziommer, D. Kilian, T. Wollborn, U. Fritsching, J. Steingroewer, T. Walther, M. Gelinsky, A. Lode, Biofabrication 2020, 12, 025022; bR. V. Barrulas, M. C. Corvo, Gels 2023, 9.

[14] Y. P. Wang, Y. Z. Chen, J. N. Zheng, L. R. Liu, Q. Q. Zhang, Acs Omega 2022, 7, 12076.

[15] aW. Li, D. Wang, B. Chen, K. Hua, Z. Huang, C. Xiong, H. Yu, Materials (Basel) 2022, 15; bS. Jeon, S. H. Lee, S. B. Ahmed, J. Han, S. J. Heo, H. W. Kang, Essays Biochem 2021, 65, 467; cM. A. Skylar-Scott, S. G. M. Uzel, L. L. Nam, J. H. Ahrens, R. L. Truby, S. Damaraju, J. A. Lewis, Sci Adv 2019, 5, eaaw2459.

[16] S. Bhumiratana, J. Bernhard, E. Cimetta, G. Vunjak-Novakovic, in Principles of Tissue Engineering, 2014, 261.

