## Supplemental figures for "Biofabrication of engineered tissues by 3D bioprinting of tissue specific high cell-density bioinks"

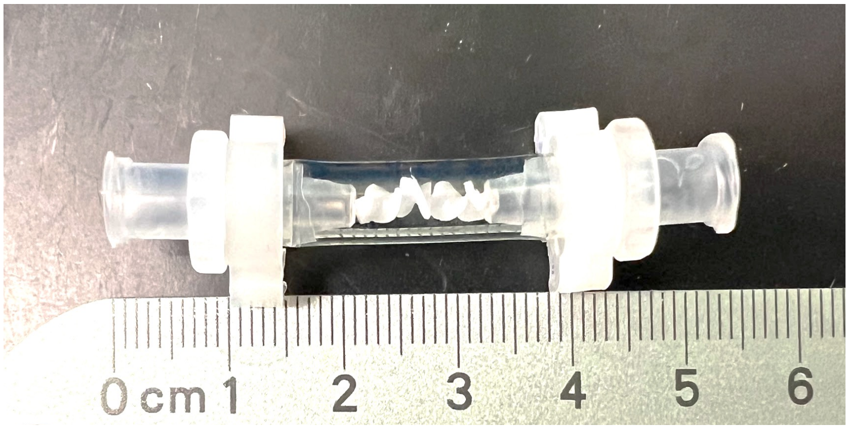


**Figure S1.** Photograph of the custom spiral mixing unit.

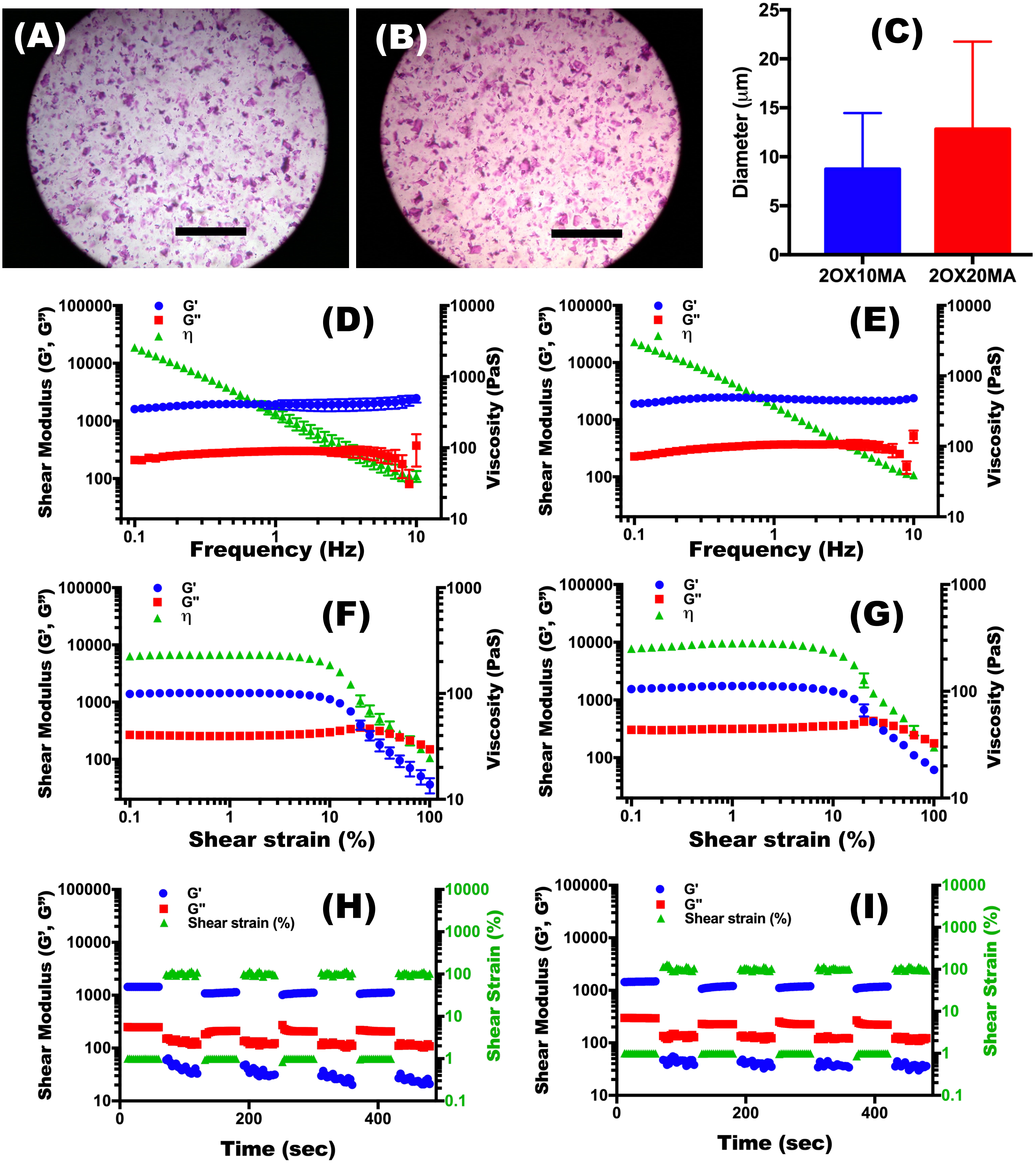


**Figure S2.** Optical photomicrograph of Safranin-O stained (A) 2OX10MA and (B) 2OX20MA OMA microgels and (C) their mean diameters. Scale bars indicate 1 mm. Storage modulus (G'), loss modulus (G") and viscosity (η) of (D) 2OX10MA and (E) 2OX20MA OMA microgels as a function of frequency (Hz). Storage modulus (G'), loss modulus (G") and viscosity (η) of (F) 2OX10MA and (G) 2OX20MA OMA microgels as a function of shear strain (%). Shear moduli (G' and G") changes in dynamic strain tests of the (H) 2OX10MA and (I) 2OX20MA OMA microgels with alternating low (1 %) and high (100 %) strains at 1 Hz frequency.

(A)

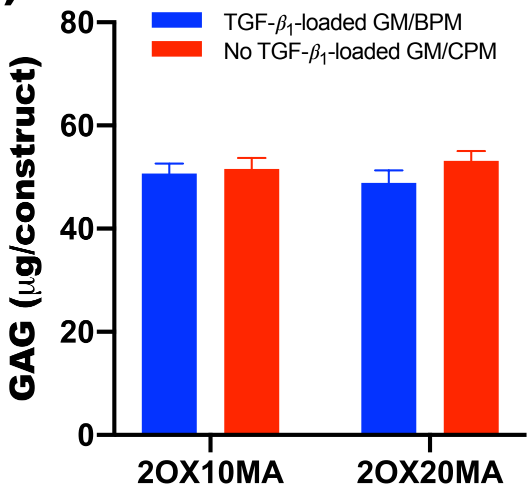


(B)

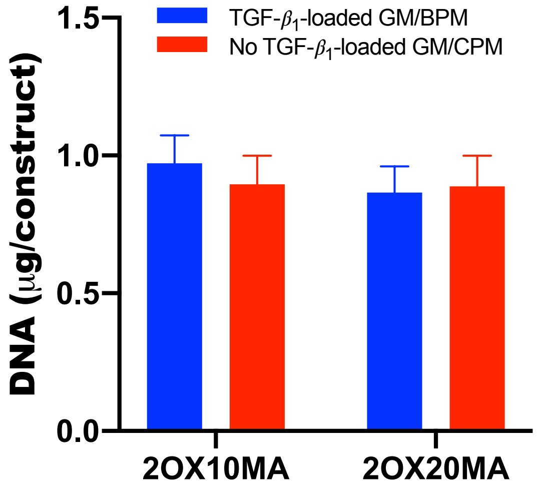


**Figure S3.** Quantification of (A) GAG production and (B) DNA content in the 3D printed constructs after 4 weeks of culture.

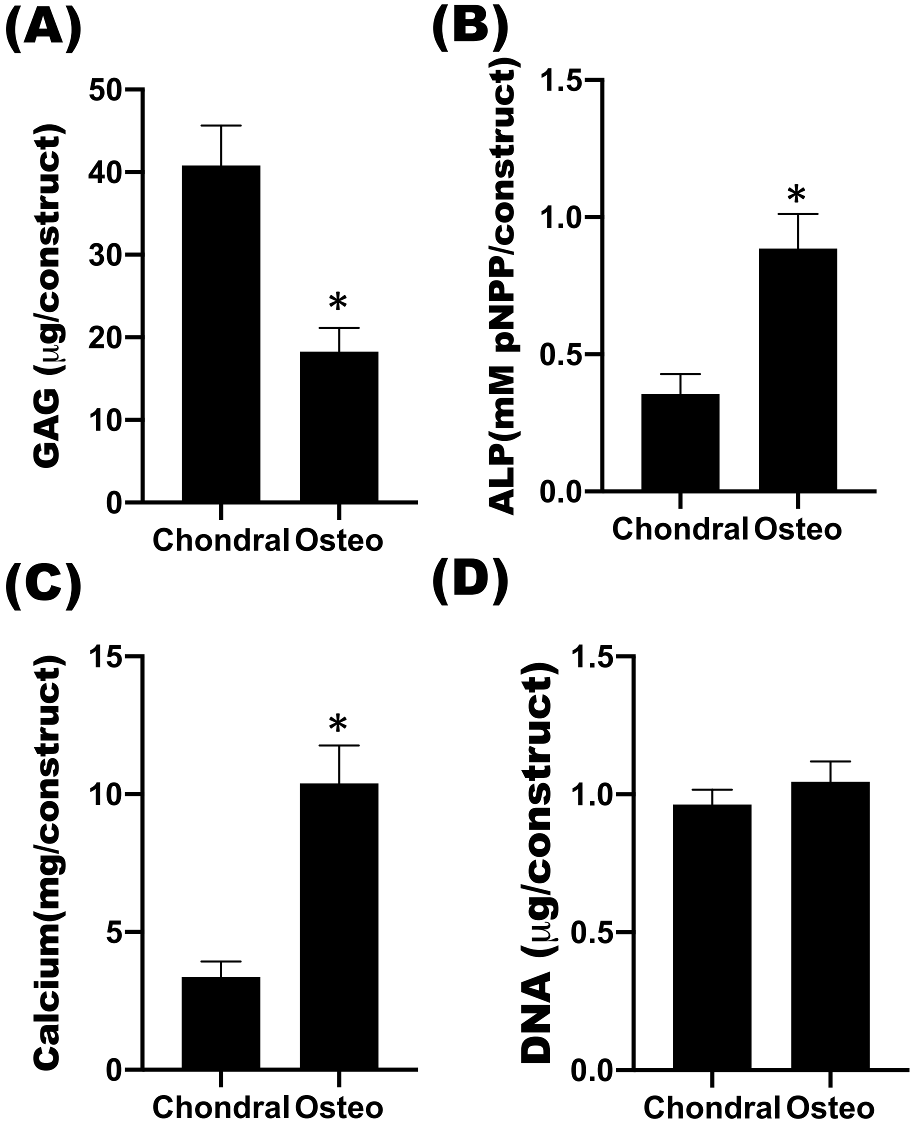


**Figure S4.** Quantification (A) GAG production, (B) ALP activity, (C) calcium content and (D) DNA content of chondrogenic (Chondral) and osteogenic (Osteo) phases of the osteochondral constructs. *$p<0.05$ compared to chondrogenic phase.

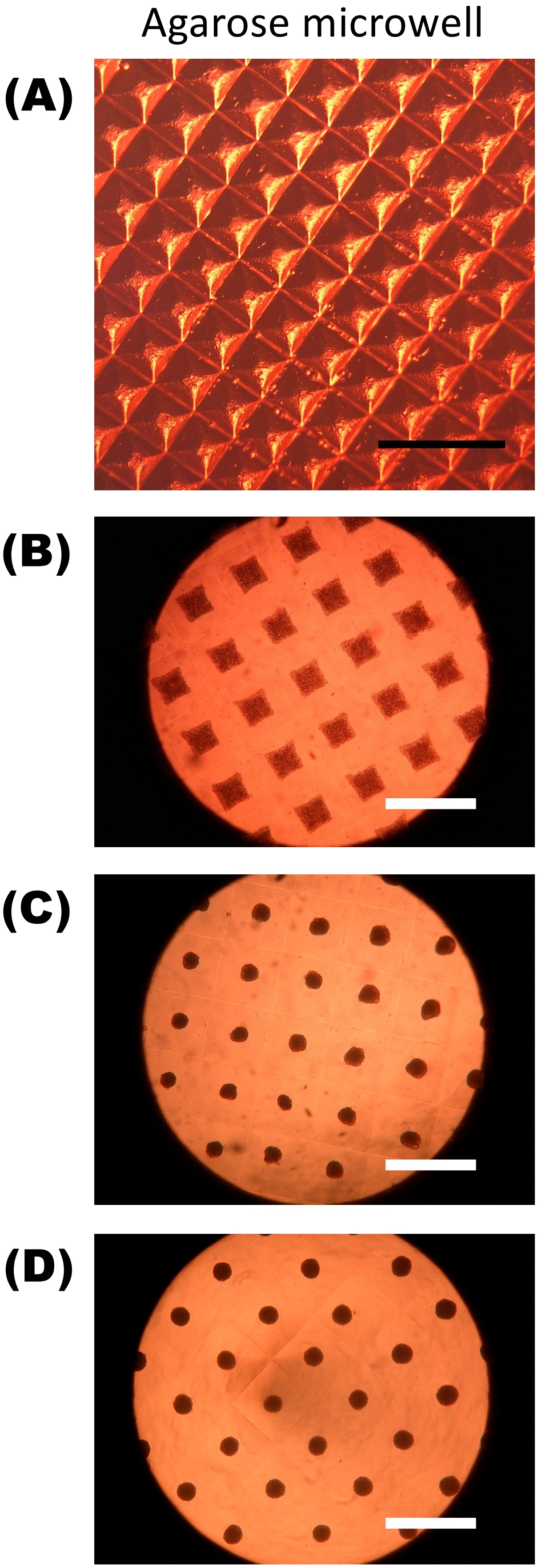


**Figure S5.** (A) Optical photomicrograph of agarose microwells. Scale bar indicates 2 mm. Optical photomicrographs of hMSCs in the agarose microwells at (B) Day 0, (C) Day 1 and (D) Day 2. Scale bars indicate 500 μm.

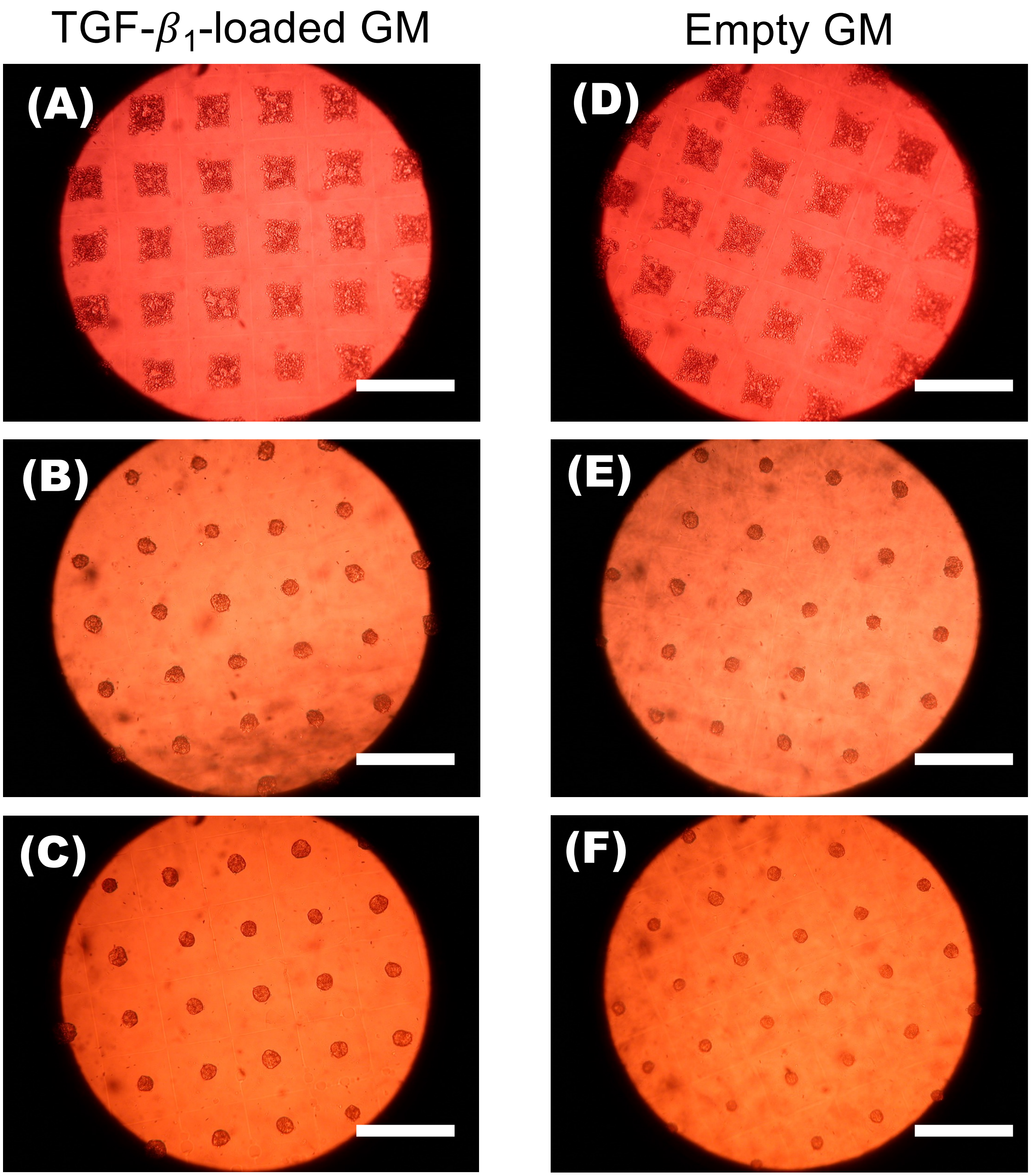


**Figure S6.** Optical photomicrographs of hMSCs/TGF-$\beta_1$-loaded GM in the agarose microwells at (A) Day 0, (B) Day 1 and (C) Day 2. Optical photomicrographs of hMSCs/Empty GM in the agarose microwells at (D) Day 0, (E) Day 1 and (F) Day 2. Scale bars indicate 500 μm.

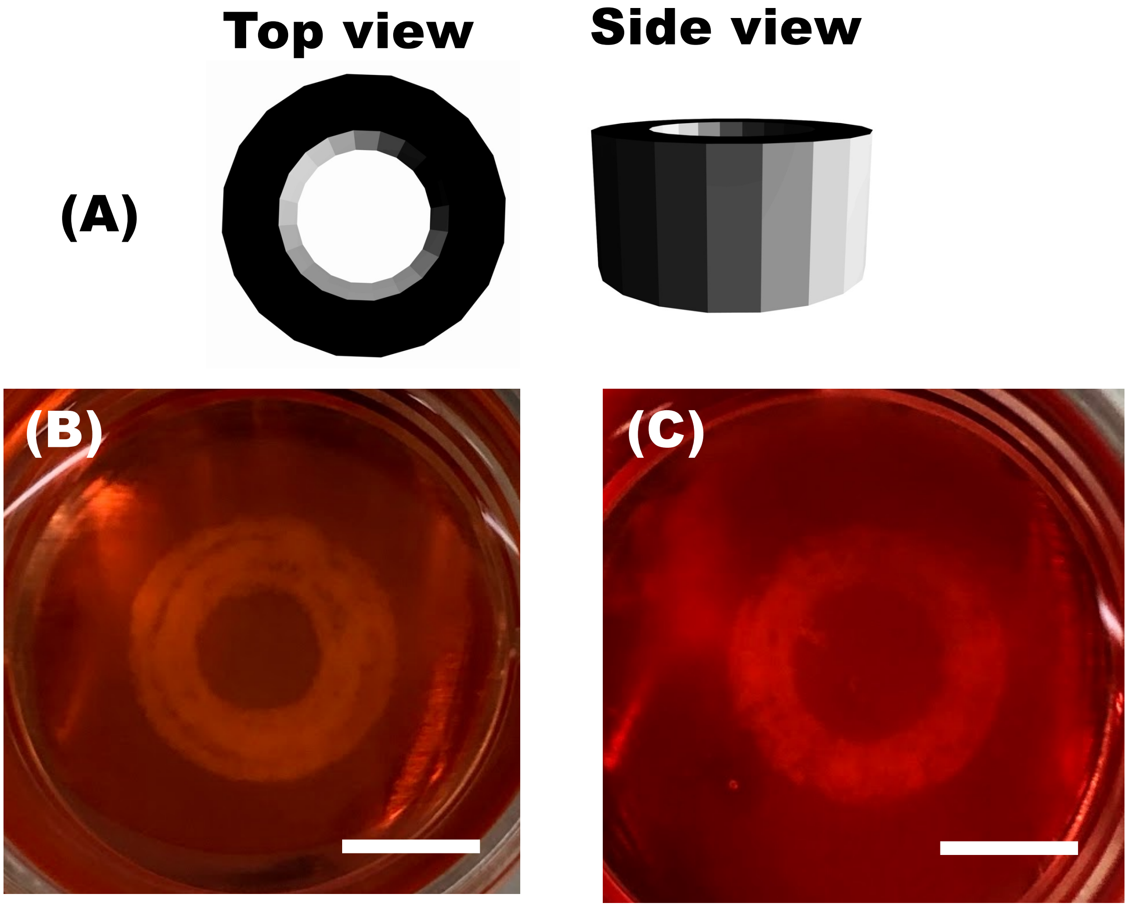


**Figure S7.** (A) Digital images and photographs of 3D printed tubes in the photocrosslinkable alginate microgel slurry bath using (B) aggregates and GM mixed and (C) aggregates formed with GM bioinks. Scale bars indicate 5 mm.

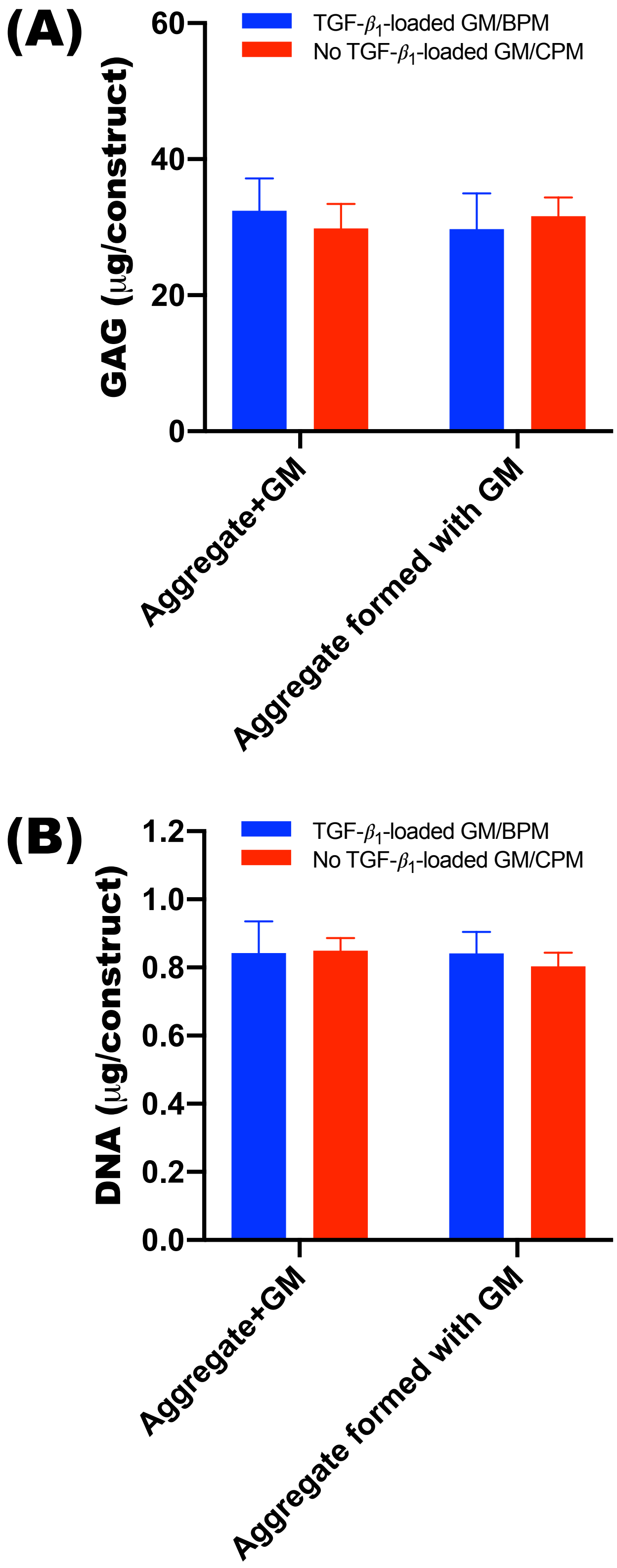


**Figure S8.** Quantification of (A) GAG production and (B) DNA content in the 3D printed constructs after 4 weeks culture.
